# SARS-CoV-2 and SARS-CoV differ in their cell tropism and drug sensitivity profiles

**DOI:** 10.1101/2020.04.03.024257

**Authors:** Denisa Bojkova, Jake E. McGreig, Katie-May McLaughlin, Stuart G. Masterson, Marek Widera, Verena Krähling, Sandra Ciesek, Mark N. Wass, Martin Michaelis, Jindrich Cinatl

## Abstract

SARS-CoV-2 is a novel coronavirus currently causing a pandemic. We show that the majority of amino acid positions, which differ between SARS-CoV-2 and the closely related SARS-CoV, are differentially conserved suggesting differences in biological behaviour. In agreement, novel cell culture models revealed differences between the tropism of SARS-CoV-2 and SARS-CoV. Moreover, cellular ACE2 (SARS-CoV-2 receptor) and TMPRSS2 (enables virus entry via S protein cleavage) levels did not reliably indicate cell susceptibility to SARS-CoV-2. SARS-CoV-2 and SARS-CoV further differed in their drug sensitivity profiles. Thus, only drug testing using SARS-CoV-2 reliably identifies therapy candidates. Therapeutic concentrations of the approved protease inhibitor aprotinin displayed anti-SARS-CoV-2 activity. The efficacy of aprotinin and of remdesivir (currently under clinical investigation against SARS-CoV-2) were further enhanced by therapeutic concentrations of the proton pump inhibitor omeprazole (aprotinin 2.7-fold, remdesivir 10-fold). Hence, our study has also identified anti-SARS-CoV-2 therapy candidates that can be readily tested in patients.

## Introduction

In December 2019, SARS-CoV-2, a novel betacoronavirus, was identified that causes a respiratory disease and pneumonia called coronavirus disease 19 (COVID-19) [Chen et al., 2020; Coronaviridae Study Group of the International Committee on Taxonomy of Viruses, 2020; Lu et al., 2020; Wu et al., 2020; Zhou et al., 2020; Zhu et al., 2020]. The first cases seemed to have originated from a wholesale fish market in Wuhan, China [Zhu et al., 2020]. As of 3^rd^ April 2020, this novel virus has resulted in 1,041,126 confirmed infections and 55,132 deaths in 181 countries and regions (www.who.int) [Dong et al., 2020].

SARS-CoV-2 is the seventh coronavirus known to infect and cause disease in humans alongside the alphacoronaviruses human coronavirus 229E (HCoV-229E) and human coronavirus NL63 (HCoV-NL63, New Haven coronavirus) and the betacoronaviruses human coronavirus OC43 (HCoV-OC43), human coronavirus HKU1 (HCoV-HKU1), severe acute respiratory syndrome coronavirus (SARS-CoV), and Middle East respiratory syndrome coronavirus (MERS-CoV) [Corman et al., 2018; Yin and Wunderink, 2018; Cui et al., 2019; Wu et al., 2020b]. HCoV-229E, HCoV-OC43, HCoV-NL63, and HCoV-HKU1 are endemic in humans and typically cause mild to moderate common cold-like respiratory disease [Channappanavar & Perlman, 2017; Corman et al., 2018].

Since 2002, SARS-CoV-2 is the third coronavirus, after SARS-CoV and MERS-CoV, that has caused a substantial outbreak associated with significant mortality [Wu et al., 2020b]. According to WHO, the SARS-CoV outbreak resulted in 8,098 confirmed and suspected cases and 774 deaths, equalling a mortality rate of 9.6% (www.who.int). For MERS-CoV, the WHO currently (2nd April 2020) reports 2,494 laboratory-confirmed cases and 858 deaths (mortality rate: 34.4%) (www.who.int). However, human-to-human spread of MERS-CoV remains very limited. SARS-CoV-2 disease is associated with a lower mortality. Currently, about 5.3% of individuals with confirmed SARS-CoV-2 infection have died, and the risk of severe disease increases with age [Dong et al., 2020; CDC COVID-19 Response Team, 2020]. This mortality level is likely to be an overestimation. A mortality rate of 1% or less may be more realistic, because patients with severe symptoms are more likely to be tested, while mild and asymptomatic cases are likely to go unreported [Borges do Nascimento, 2020; Nishiura et al., 2020; Pan et al., 2020; Rothe et al., 2020]. In contrast to SARS-CoV-infected patients, SARS-CoV-2 has been reported to be spread by individuals who are asymptomatic during the incubation period or who do not develop symptoms at all [Li et al., 2020; Nishiura et al., 2020; Nishiura et al., 2020a; Pan et al., 2020; Rothe et al., 2020; Yu et al., 2020]

We have developed an approach to identify sequence-associated phenotypic differences between related viruses based on the identification of differentially conserved amino acid sequence positions (DCPs) and in silico modelling of protein structures [Pappalardo et al., 2016; Martell et al., 2019]. Here, we used this method to identify differentially conserved positions that may explain phenotypic differences between SARS-CoV-2 and SARS-CoV [Coronaviridae Study Group of the International Committee on Taxonomy of Viruses, 2020; Lu et al., 2020; Zhou et al., 2020]. These findings were analysed in combination with data from cells infected with a recently derived SARS-CoV-2 isolate [Hoehl et al., 2020]. Our results reveal characteristic differences between SARS-CoV-2 and SARS-CoV. Most importantly, we found that therapeutic concentrations of the protease inhibitor aprotinin interfere with SARS-CoV-2 infection. The efficacy of aprotinin can be further increased by therapeutic concentrations of the proton pump inhibitor omeprazole.

## Results

### Determination of differentially conserved positions (DCPs)

Coronavirus genomes harbour single-stranded positive sense RNA (+ssRNA) of about 30 kilobases in length, which contain six or more open reading frames (ORFs) [Cui et al., 2019; Song et al., 2019; Chen et al., 2020b; Wu et al., 2020b]. The SARS-CoV-2 genome has a size of approximately 29.8 kilobases and was annotated to encode 14 ORFs and 27 proteins [Wu et al., 2020b]. Two ORFs at the 5’-terminus (ORF1a, ORF1ab) encode the polyproteins pp1a and pp1b, which comprise 15 non-structural proteins (nsps), the nsps 1 to 10 and 12-16 [Wu et al., 2020b]. Additionally, SARS-CoV-2 encodes four structural proteins (S, E, M, N) and eight accessory proteins (3a, 3b, p6, 7a, 7b, 8b, 9b, orf14) [Wu et al., 2020b]. This set-up resembles that of SARS-CoV. Notable differences include that there is an 8a protein in SARS-CoV, which is absent in SARS-CoV-2, that 8b is longer in SARS-CoV-2 (121 amino acids) than in SARS-CoV (84 amino acids), and that 3b is shorter in SARS-CoV-2 (22 amino acids) than in SARS-CoV (154 amino acids) [Wu et al., 2020b].

To identify genomic differences between SARS-CoV-2 and SARS-CoV that may affect the structure and function of the encoded virus proteins, we applied an approach that we have previously used to compare the human pathogenic Ebolaviruses species with Reston virus, an Ebolavirus that does not cause disease in humans [Pappalardo et al., 2016; Martell et al., 2019]. This methodology is based on the determination of differentially conserved positions (DCPs) [Rausell et al., 2010], i.e. amino acid positions that are differently conserved between phenotypically different groups, in our case related viruses. The potential impact of the DCPs on protein structure and function is then determined by in silico modelling [Pappalardo et al., 2016; Martell et al., 2019].

For the 22 SARS-CoV-2 virus proteins that could be compared with SARS-CoV, comparison of the two reference sequences identified 1393 positions that encode different amino acids. 1243 (89%) of these positions were DCPs (Table 1), which represents 13% of all residues encoded by the SARS-CoV-2 genome. Most of the amino acid substitutions at DCPs appear to be fairly conservative as demonstrated by the average BLOSUM substitution score of 0.49 (median 0; Supplementary Figure 1) and with 73% of them having a score of 0 or greater (the higher the score the more frequently such amino acid substitutions are observed naturally in evolution). It followed that 45% of DCPs represent conservative changes where amino acid properties are retained (e.g. change between two hydrophobic amino acids), a further 30% represented polar - hydrophobic substitutions, while changes between charged amino acids were rare (<10% of DCPs) (Supplementary Table 1).

**Table 1.**
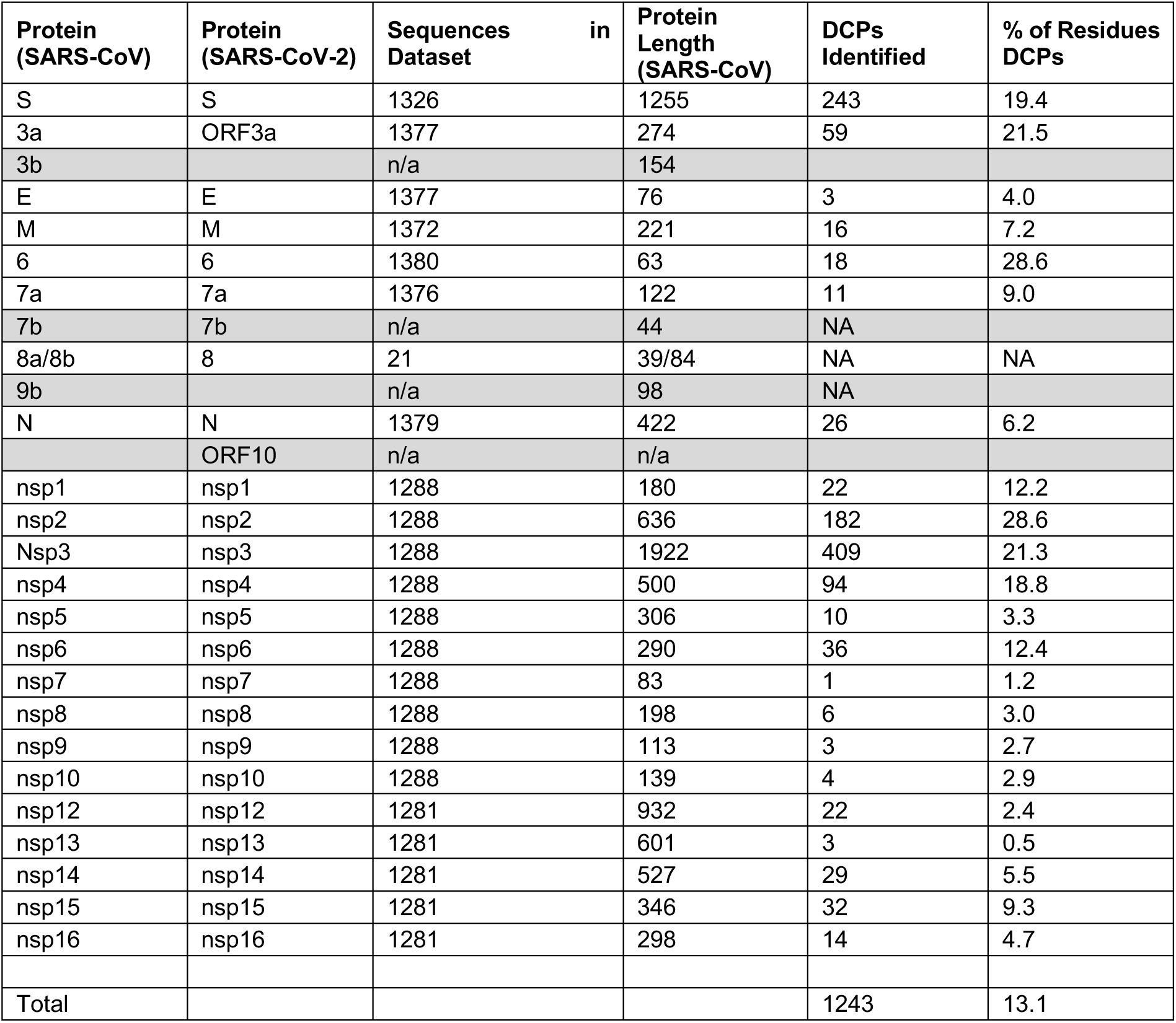
Specificity Determining Positions (DCPs) identified between SARS-CoV and SARS-CoV-2.

DCPs are enriched in six of the SARS-CoV-2 proteins, spike (S), 3a, p6, nsp2, nsp3 (papain-like protease) and nsp4 with 19.4%, 21.5%, 28.6%, 28.6%, 21.3% and 18.8% of their residues being DCPs respectively (Table 1). In contrast, very few DCPs were observed in the envelope (E) protein and most of remaining non-structural proteins encoded by ORF1ab, for example 0.5% of residues in the helicase and 2% of residues in the RNA-directed RNA polymerase, 2’-O-Methyltransferase, nsp8 and nsp9 are DCPs (Table 1).

The availability of structures of both SARS-CoV and some SARS-CoV-2 proteins, coupled with the ability to model some of the remaining proteins (ref to methods and supplementary table) enabled us to map 525 DCPs onto protein structures (Supplementary Figure 1, Supplementary Table 1). Overall, nearly all of the mapped DCPs occur on the protein surface (92%), with only 40 DCPs buried within the protein, primarily in S and the papain-like protease (nsp3) (Supplementary Table 1). Based on our structural analysis, we propose that 45 DCPs are likely to result in structural (or functional) differences between SARS-CoV and SARS-CoV-2 proteins. A further 222 could result in some change, with our analysis suggesting that the remaining 258 DCPs seem unlikely to have a substantial effect on protein structure and function.

### Differentially conserved positions (DCPs) in interferon antagonists

At least 10 SARS-CoV proteins have roles in interferon antagonism [Totura and Baric, 2012]. Two of these proteins, p6 and the papain-like protease (nsp3), are enriched in DCPs, two are depleted in DCPs (nsp7 and nsp16), five have intermediate proportions of DCPs (nsp14, nsp1, nsp15, N and M), while p3b is not encoded by SARS-CoV-2. Initial studies have identified a difference in the interferon inhibition between SARS-CoV and SARS-CoV-2 [Lokugamage et al., 2020], so it is possible that the DCPs identified in these proteins, especially in p6 and the papain-like protease, may have an effect on interferon inhibition.

### S (Spike) protein

The most interesting changes were detected in the spike (S) protein, which mediates coronavirus entry into host cells [Cui et al., 2019; Chen et al., 2020b]. SARS-CoV-2 S is 77.46% sequence identical to the SARS-CoV S and most of the remaining positions are DCPs (243 residues, 1%) (Table 1). SARS-CoV entry depends on the cleavage of S by transmembrane serine protease 2 (TMPRSS2), the endosomal cysteine protease cathepsin L, and/ or other cellular proteases, with residues R667 and R797 being critical cleavage sites [Matsuyama et al., 2010; Simmons et al., 2013; Zhou et al., 2015; Reinke et al., 2017; Iwata-Yoshikawa et al., 2019]. Serine protease inhibitors such as camostat and nafamostat interfere with S cleavage by TMPRSS2 and virus uptake [Kawase et al., 2012; Yamamoto et al., 2016; Zhou et al., 2015; Shin & Seong, 2017]. R667 and R797 are conserved in SARS-CoV-2 (R685 and R815). However, there is a four amino acid insertion in SARS-CoV-2 S prior to R685 and many of the residues close to R685 are DCPs (V663=Q677, S664=T678, T669=V687, S670=A688, Q671=S689, DCPs are represented by the SARS CoV residue followed by the SARS-CoV-2 residue) (Figure 1A). There is greater conservation around the R815 cleavage site with only two DCPs in close proximity (L792=S810, T795=S813) (Figure 1A).

**Figure 1.**
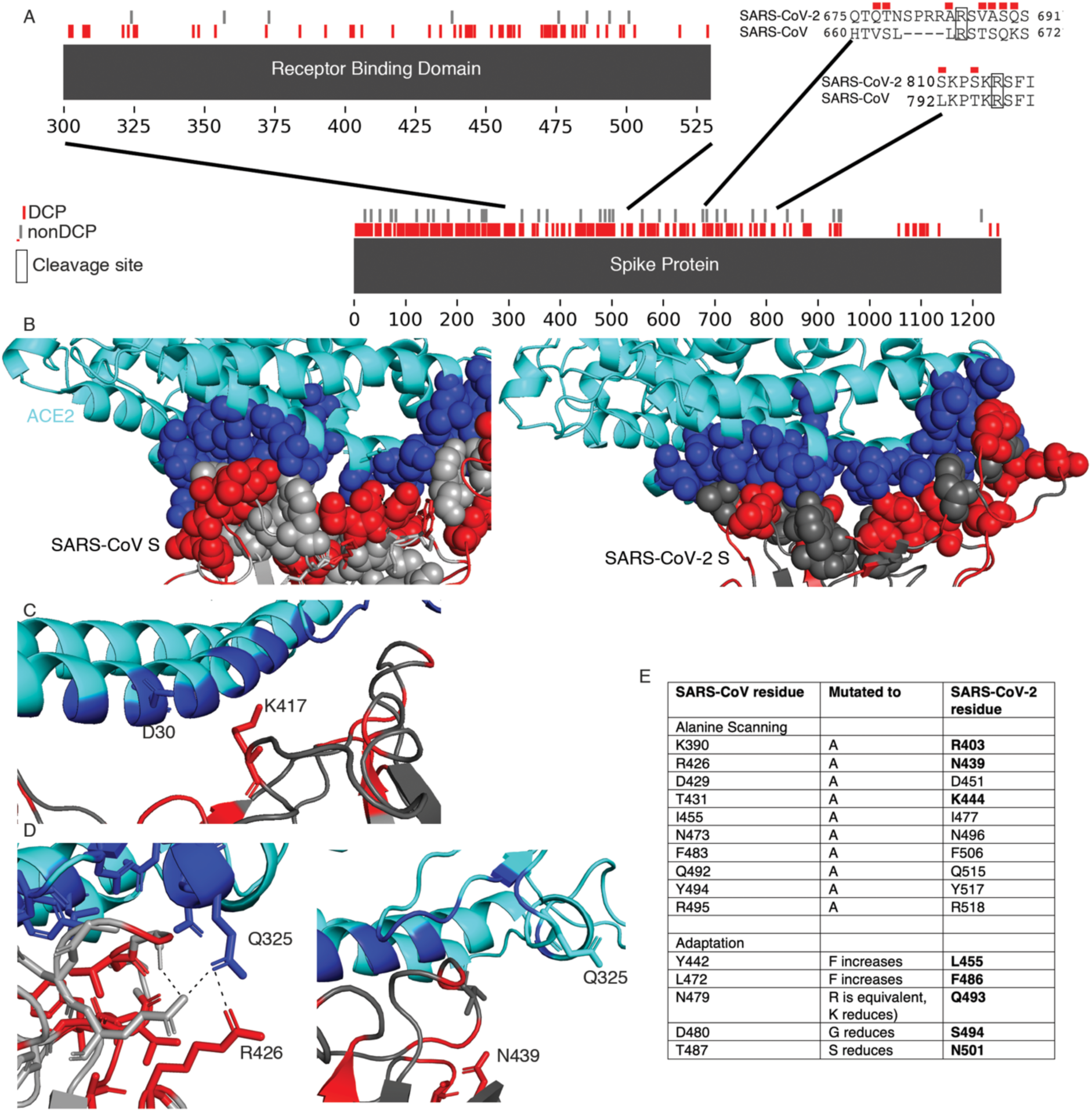
Differentially conserved positions in the Spike protein. A) A sequence view of the DCPs present in the Spike protein, with insets showing the receptor binding domain and the two cleavage sites. B) The S interface with ACE2 (cyan). The ACE2 interface is shown in blue spheres, DCPs in red. C) The V404=K417 DCP. D) The R426=N439 DCP, the left image shows SARS-CoV S R426, the image on the right show the equivalen N439 in SARS-CoV-2 S. E) SARS-CoV residues associated with altering ACE2 affinity and the residues at these positions in SARS-CoV-2 S.

The SARS-CoV S receptor binding domain (residues 306-527, equivalent to 328-550 in SARS-CoV-2) is enriched in DCPs, containing 51 DCPs (23% of residues). Eleven of the 24 SARS-CoV S residues in direct contact with ACE2 were DCPs (Figure 1A, Supplementary Table 2). Analysis of the DCPs using the SARS-CoV and SARS-CoV-2 S protein complexes with ACE2 [Song et al., 2018; Yan et al., 2020] identified runs of DCPs (A430-T433, F460-A471) in surface loops forming part of the S-ACE2 interface and resulted in different conformations in SARS-CoV-2 S compared to SARS-CoV S (Figure 1B). Two DCPs remove intramolecular hydrogen bonding within the spike protein in SARS-CoV-2 (Supplementary Table 2) and three DCPs (R426=N439, N479=QQ493, Y484=Q498) are residues that form hydrogen bonds with ACE2. For two of these positions, hydrogen bonding with ACE2 is present with both S proteins, but for R426=N439 hydrogen bonding with ACE2 is only observed with SARS-CoV S. N439 in SARS-CoV-2 S is not present in the interface and the sidechain points away from the interface (Figure 1D). Further, analysis of the SARS-CoV-2 S-ACE2 complex highlighted important roles of the V404=K417 DCP, where K417 in SARS-CoV-2 S is able to form a salt bridge with ACE2 D30 (Figure 1C) [Yan et al., 2020].

Alanine scanning [Chakraborti et al., 2005] and adaptation experiments [Wan et al., 2020] have identified 16 SARS-CoV S residues associated with determining the binding affinity with ACE2. For all five residues identified from adaptation studies and four of the 11 identified by alanine scanning epxeriments different amino acids are present in SARS-CoV-2 S (Figure 1E), highlighting the difference in the interaction with ACE2.

### SARS-CoV-2 replication in different cell lines

In further experiments, we investigated to see the extent to which the substantial number of amino acid positions that are differently conserved between SARS-CoV and SARS-CoV-2, result in different phenotypes. Infection experiments using the four colorectal cancer cell lines Caco2, CL14, HT-29, and DLD-1 resulted in similar susceptibility profiles. Replication of both viruses was detected in Caco2 and CL14 cells, but not in HT-29 or DLD-1 cells, as shown by cytopathogenic effects (CPE) (Figure 2A), staining for double-stranded RNA (Figure 2B), and viral genomic RNA levels (Figure 2C). These findings are in line with previous findings showing that Caco2 and CL14 cells are susceptible to SARS-CoV infection [Cinatl et al., 2004] and the previous isolation of SARS-CoV-2 cells in Caco2 cells [Hoehl et al., 2020]. Moreover, we identified CL14 as an additional model to study SARS-CoV-2 infection and replication.

**Figure 2.**
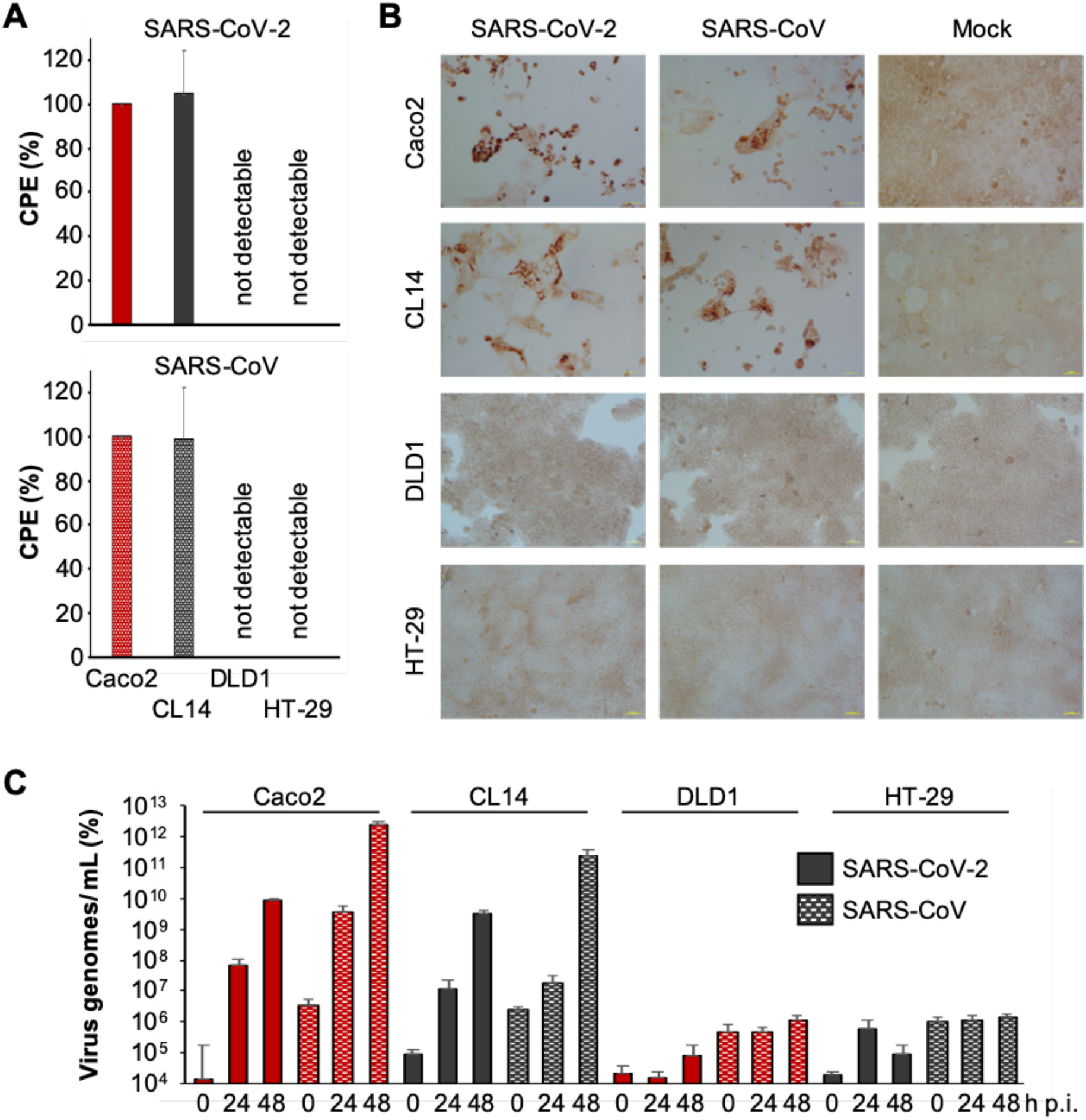
SARS-CoV-2 and SARS-CoV susceptibility of colorectal cancer cell lines. A) Cytopathogenic effect (CPE) formation 48h post infection in MOI 0.01-infected cells. B) Representative images showing MOI 0.01-infected cells immunostained for double-stranded RNA 48h post infection. C) Quantification of virus genomes by qPCR at different time points post infection (p.i.).

### SARS-CoV-2 infection does not correlate with the cellular ACE2 status

Although ACE2 was identified as a SARS-CoV-2 receptor [Hoffmann et al., 2020; Letko et al., 2020; Walls et al., 2020; Wan et al., 2020; Wrapp et al., 2020; Yan et al., 2020; Zhou et al., 2020], there was no correlation between the cellular ACE2 levels and the cellular susceptibility to SARS-CoV-2 (and SARS-CoV) (Figure 3A). CL14 cells displayed lower ACE2 levels than both HT-29 and DLD-1 (Figure 3A), although CL14 was, in contrast to HT-29 and DLD-1, permissive to SARS-CoV-2 (and SARS-CoV) infection (Figure 2). This suggests that there are other factors in addition to the cellular ACE2 levels that determine cellular susceptibility to SARS-CoV-2 and SARS-CoV.

**Figure 3.**
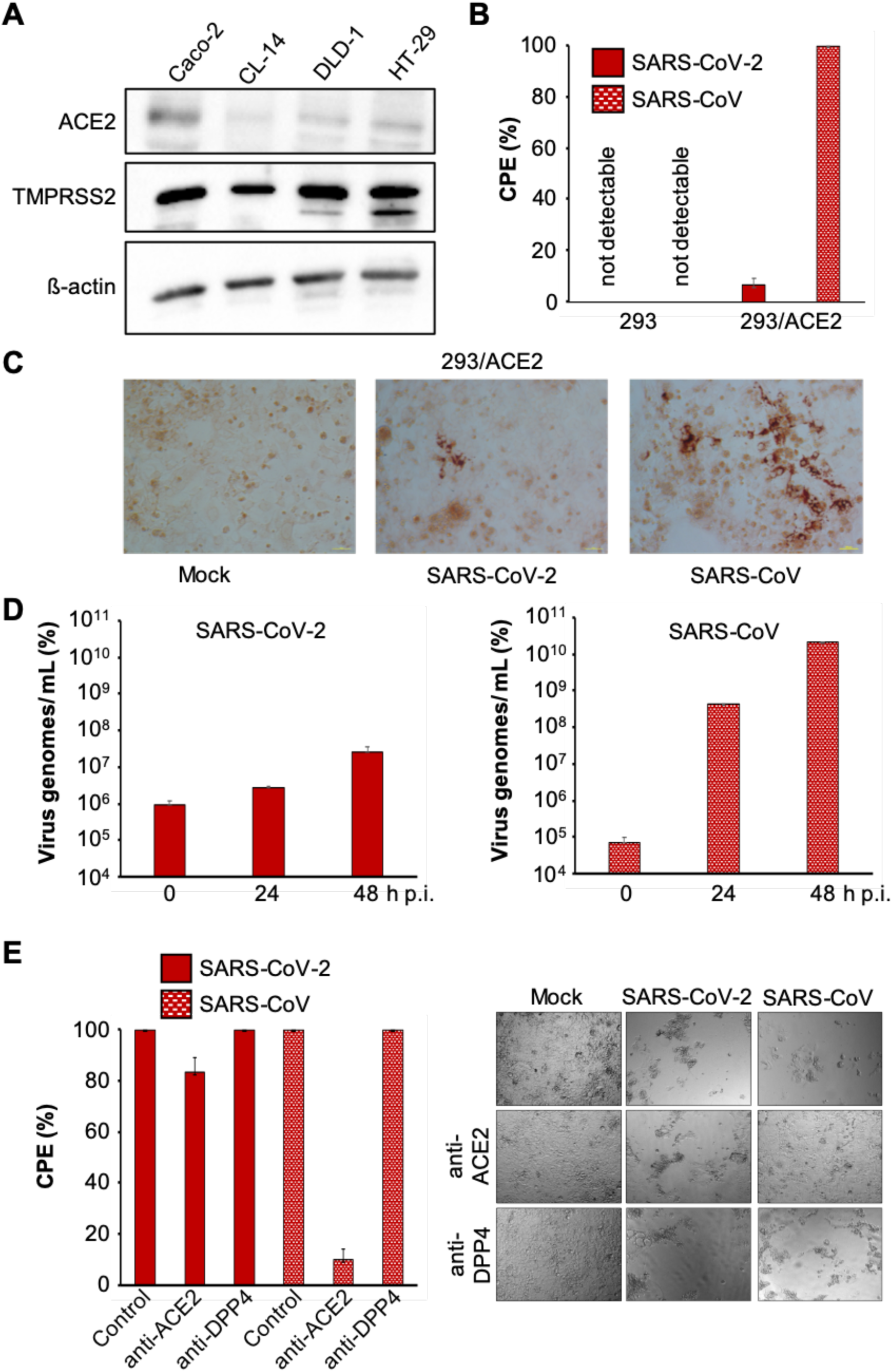
N A) Western blots indicating cellular ACE2 and TMPRSS2 protein levels. B) CPE formation in SARS-CoV and SARS-CoV-2 (MOI 0.01)-infected ACE2-negative 293 cells and 293 cells stably expressing ACE2 cells (293/ACE2) 48h post infection. C) Immunostaining for double-stranded RNA in SARS-CoV-2 and SARS-CoV (MOI 0.01)-infected 293/ACE2 cells 48h post infection. D) Quantification of virus genomes by qPCR in SARS-CoV-2 and SARS-CoV (MOI 0.01)-infected 293/ACE2 cells 48h post infection. E) Cytopathogenic effect (CPE) formation in SARS-CoV-2 and SARS-CoV (MOI 0.01)-infected Caco2 cells in the presence of antibodies directed against ACE2 or DPP4 (MERS-CoV receptor) 48h post infection.

### ACE2 expression mediates 293 cell susceptibility to SARS-CoV but not to SARS-CoV-2

Next, we compared SARS-CoV-2 and SARS-CoV replication dependence on ACE2 in an additional model. 293 cells are not susceptible to SARS-CoV infection due to a lack of ACE2 expression. However, 293 cells that stably express ACE2 (293/ACE2) support SARS-CoV infection [Kamitani et al., 2006]. As expected, infection of 293 cells with SARS-CoV or SARS-CoV-2 did not result in detectable cytopathogenic effect (CPE) (Figure 3B), but a SARS-CoV-induced CPE was detected in 293/ACE2 cells (Figure 3B). In contrast to SARS-CoV, however, 293/ACE2 cells displayed limited permissiveness to SARS-CoV-2 infection (Figure 3B). Staining for double-stranded RNA (Figure 3C) and detection of viral genomic RNA copies (Figure 3D) confirmed reduced SARS-CoV-2 infection of and replication in 293/ACE2 cells relative to SARS-CoV. These findings further suggest that there are differences in the host cell factors that mediate SARS-CoV and SARS-CoV-2 susceptibility and in turn differences in the cell tropisms.

### Reduced activity of anti-ACE2 antibody against SARS-CoV-2 compared to SARS-CoV

Antibodies directed against ACE2 have been shown to inhibit SARS-CoV replication [Li et al., 2003]. In agreement, an anti-ACE2 antibody inhibited SARS-CoV infection in Caco2 cells (Figure 3E). However, the anti-ACE2 antibody displayed limited activity against SARS-CoV-2 infection (Figure 3E). This is in agreement with previous findings indicating a stronger binding affinity of SARS-CoV-2 S to ACE2 compared to SARS-CoV S [Walls et al., 2020; Wrapp et al., 2020], which may be more difficult to antagonise using anti-ACE2 antibodies. As anticipated, antibodies directed against DPP4, the MERS-CoV receptor [de Wit et al., 2016; Cui et al., 2019], did not interfere with SARS-CoV or SARS-CoV-2 infection (Figure 3E).

### SARS-CoV-2 is more sensitive to TMPRSS2 inhibitors than SARS-CoV

SARS-CoV S and SARS-CoV-2 S are cleaved and activated by TMPRSS2 (transmembrane serine protease 2) [Matsuyama et al., 2010; Hoffmann et al., 2020; Matsuyama et al., 2020]. Notably, all four cell lines, which we had tested for susceptibility to SARS-CoV-2 replication, displayed similar TMPRSS2 levels (Figure 3A). Hence, cellular permissiveness to SARS-CoV-2 infection is determined by further host cell factors in addition to TMPRSS2 and ACE2.

Previous findings had shown that the serine protease inhibitor camostat, which is approved for the treatment of chronic pancreatitis in Japan [Ramsey et al., 2019], inhibits both SARS-CoV and SARS-CoV-2 cell entry via interference with TMPRSS2-mediated S cleavage [Kawase et al., 2012; Zhou et al., 2015; Hoffmann et al., 2020]. Camostat inhibited cell entry of VSV pseudotypes bearing SARS-CoV-2 S in a concentration-dependent manner [Hoffmann et al., 2020]. Control experiments using wild-type virus were only performed using a single high camostat concentration of 100 µM. Here, we directly compared the concentration-dependent effects of camostat on SARS-CoV-2- and SARS-CoV-induced CPE formation in Caco2 cells (Figure 4A). Camostat displayed nearly 14-fold increased activity against SARS-CoV-2 (concentration that inhibits CPE formation by 50%, IC_50_ 1.20µM) compared to SARS-CoV (IC_50_ 16.7µM) (Figure 4A). Nafamostat is an alternative serine protease inhibitor, which is approved for pancreatitis and [Minakata et al., 2019; Hirota et al., 2020] and has been shown to exert superior effects against MERS-CoV compared to camostat [Yamamoto et al., 2016].

**Figure 4.**
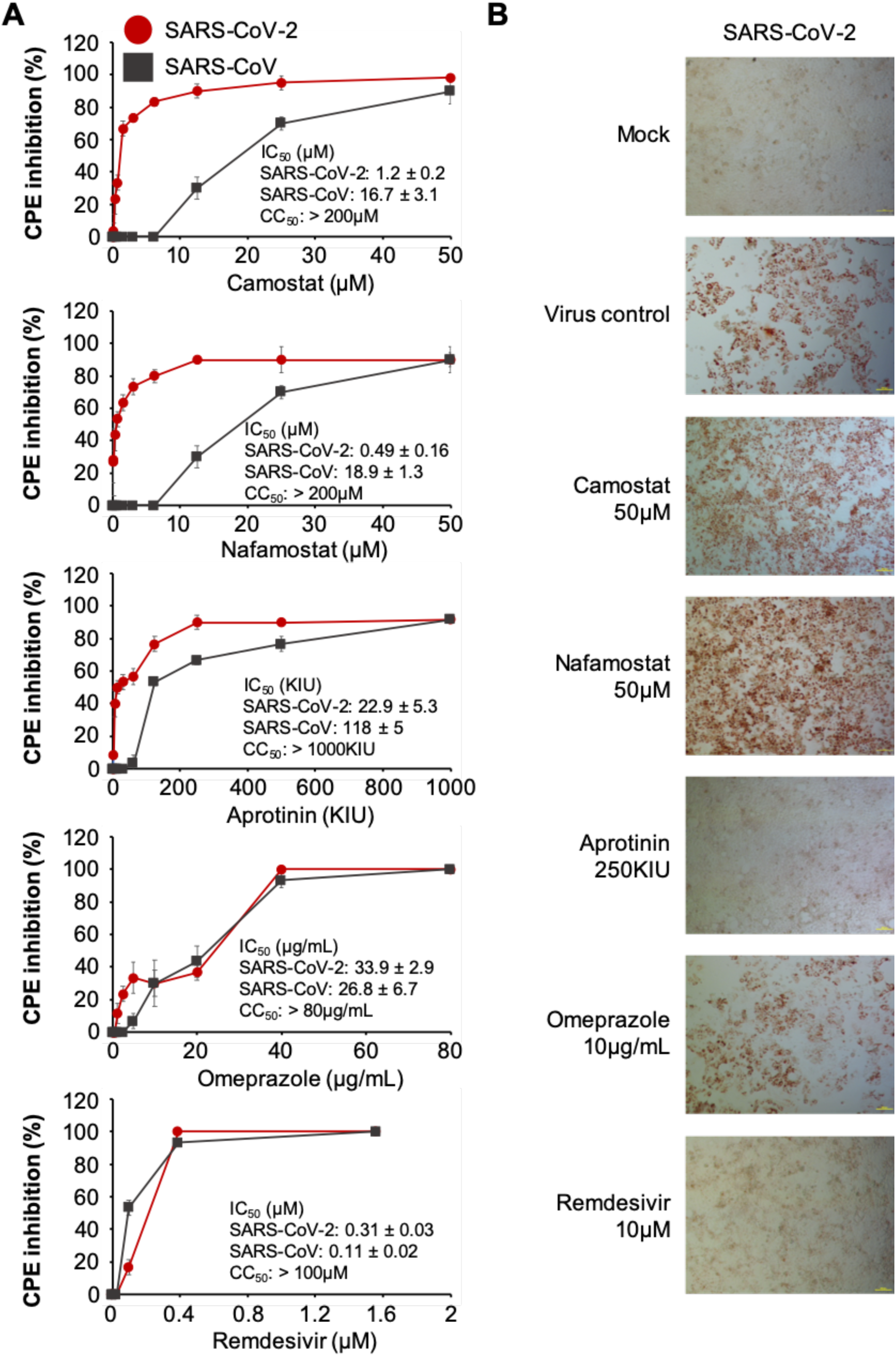
Anti-SARS-CoV-2 effects of antiviral drug candidates. A) Concentration-dependent effects of drug candidates on SARS-CoV-2- and SARS-CoV-induced cytopathogenic effect (CPE) formation determined 48h post infection in Caco2 cells infected at an MOI of 0.01. B) Immunostaining for double-stranded RNA in drug-treated SARS-CoV-2 (MOI 0.01)-infected cells 48h post infection. Camostat and nafamostat prevent SARS-CoV-2-mediated cell lysis but are characterised by high levels of double-stranded RNA.

Nafamostat displayed higher activity against SARS-CoV-2 CPE formation (IC_50_ 0.49µM) than camostat, but similar activity against SARS-CoV (18.9µM) (Figure 4A). Therapeutic plasma levels for both compounds were described to reach about 0.2µM [Hiraku et al., 1982; Cao et al., 2008], which is below the antivirally active concentrations. Moreover, although both compounds inhibited SARS-CoV-2-induced CPE formation, they displayed limited effects on the SARS-CoV-2 replication cycle as indicated by high levels of double-stranded RNA in both nafamostat- and camostat-treated SARS-CoV-2-infected cells (Figure 4B). Hence, both camostat and nafamostat may primarily exert cytoprotective effects in SARS-CoV-2-infected cells, which inhibit syncytium formation and cell lysis, but may not inhibit SARS-CoV-2 replication in the same way.

Aprotinin is a further serine protease inhibitor that has been previously investigated against influenza viruses [Zhirnov et al., 2011; Shen et al., 2017]. It is used to reduce blood loss during surgery and for pancreatitis [Moggia et al., 2017; Kapadia et al., 2019]. The efficacy of aprotinin is measured in kallikrein inhibitor units (KIU) [Levy et al., 1994; Zhirnov et al., 2011]. Like the other seine protease inhibitors, aprotinin was also more effective against SARS-CoV-2-induced CPE formation (IC_50_ 22.9 KIU/mL) than against SARS-CoV (IC_50_ 118 KIU/mL) (Figure 4A). In addition and in contrast to nafamostat and camostat, aprotinin also inhibited double-stranded RNA formation in SARS-CoV-2-infected cells (Figure 4B). Therapeutic aprotinin plasma levels were described to reach 147 ± 61 KIU/mL after the administration of 1,000,000 KIU [Levy et al., 2019]. Moreover, an aerosol preparation of aprotinin is approved for the treatment of influenza in Russia [Zhirnov et al., 2011]. Since aprotinin interferes with SARS-CoV-2 in therapeutic concentrations and displays more pronounced direct antiviral effects than camostat and nafamostat, it seems to have a greater potential for the treatment of SARS-CoV-2-infected individuals based on our data.

### Testing of additional antiviral drug candidates

To investigate whether there are also differences in the drug sensitivity profiles of SARS-CoV-2 and SARS-CoV to other antiviral drug candidates, we tested two further compounds (Figure 4A). Previous research had shown that hydroxychloroquine and ammonium chloride interfere with SARS-CoV and SARS-CoV-2 replication, as lysosomotropic agents that increase the pH in lysosomes [Talbot and Vance, 1980; Randolph and Stollar, 1990; Touret and de Lamballerie, 2020; Wang et al., 2020; Hoffmann et al., 2020]. Proton pump inhibitors including omeprazole may also inhibit virus replication by lysosomotropic and/ or other mechanisms [Dowall et al., 2016; Strickland et al., 2017; Watanabe et al., 2020]. Thus, we included omeprazole in our study. Moreover, we tested remdesivir, a drug that was developed for the treatment of flavivirus infections and displayed activity against a range of RNA viruses [Beigel et al., 2019; Hoenen et al., 2019]. Most recently, remdesivir was found to inhibit MERS-CoV and SARS-CoV-2 and suggested as a therapy candidate for SARS-CoV-2 infection [de Wit et al., 2020; Sheahan et al., 2020; Wang et al., 2020]. Currently (as of 31^st^ March 2020), there are eight active clinical trials investigating remdesivir for SARS-CoV-2-infected individuals (www.clinicaltrials.gov). Omeprazole inhibited both viruses in similar concentrations, and SARS-CoV was more sensitive to remdesivir than to SARS-CoV-2 (Figure 4A). Both omeprazole and remdesivir also inhibited the formation of double-stranded RNA (Figure 4B)

### Omeprazole increases the anti-SARS-CoV-2 activity of remdesivir and aprotinin

Omeprazole is a well-tolerated drug and a promising candidate for drug repurposing strategies [Ikemura et al., 2017]. However, the omeprazole concentrations that interfered with SARS-CoV-2 CPE formation (IC_50_ 34µM) was beyond therapeutic omeprazole plasma concentrations reported to reach about 8µM [Shin & Kim, 2013]. We have recently shown that omeprazole increases the antiviral activity of acyclovir [Michaelis et al., 2019]. Here, we combined both aprotinin and remdesivir with a fixed omeprazole concentration of 8µM, which resulted in further increased activity against CPE formation (aprotinin 2.7-fold, remdesivir 10-fold) (Figure 5A) and double-stranded RNA formation (Figure 5B). Hence, combinations of aprotinin and remdesivir with omeprazole may represent therapy candidates for the treatment of SARS-CoV-2-associated disease.

**Figure 5.**
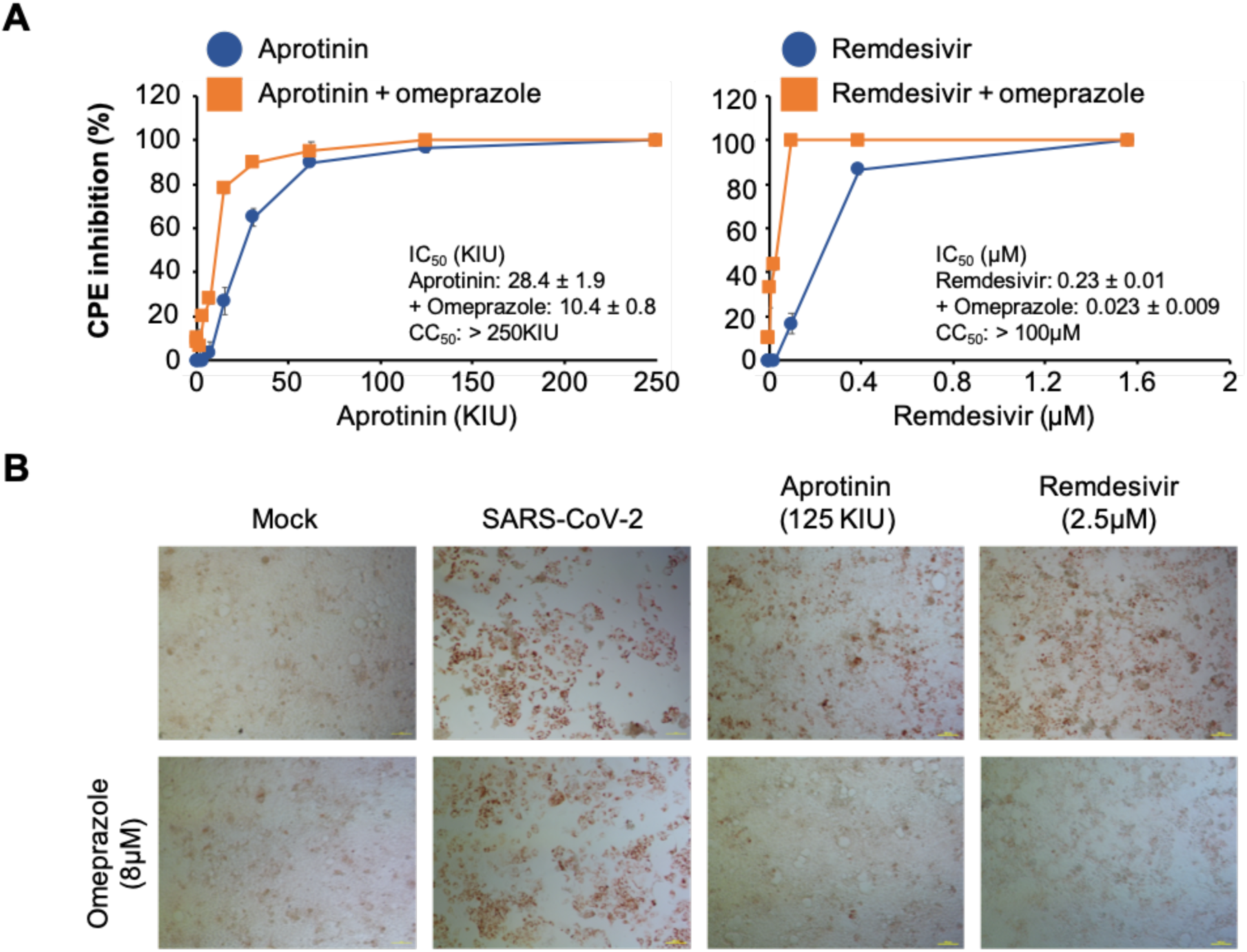
Anti-SARS-CoV-2 effects of omeprazole in combination with aprotinin and remdesivir. A) Effect of omeprazole 8µM on cytopathogenic effect (CPE) formation in combination with aprotinin and remdesivir in SARS-CoV-2 (MOI 0.01)-infected Caco2 cells 48h post infection. B) Immunostaining for double-stranded RNA indicating combined effects of aprotinin and remdesivir in combination with omeprazole in SARS-CoV-2 (MOI 0.01)-infected Caco2 cells 48h post infection.

## Discussion

Here, we performed an in-silico analysis of the effects of differentially conserved amino acid positions (DCPs) between SARS-CoV-2 and SARS-CoV proteins on virus protein structure and function in combination with a comparison of wild-type SARS-CoV-2 and SARS-CoV in cell culture.

Our analysis identified 1243 DCPs, which represents 89% of the amino acid positions that differ between SARS-CoV-2 and SARS-CoV and nearly 13% of all residues encoded by the SARS-CoV genome. 258 of these DCPs (2.6% of all residues) are likely to have a structural and functional impact. The DCPs are not equally distributed between the proteins. DCPs are enriched in S, 3a, p6, nsp2, papain-like protease, and nsp4, but very few DCPs are present in the envelope (E) protein and most of the remaining non-structural proteins encoded by ORF1ab. This indicates that the individual proteins differ in their tolerance to sequence changes and/ or their exposure to selection pressure exerted by the host environment.

This large proportion of DCPs reflects the differences in the clinical behaviour of SARS-CoV-2 and SARS-CoV. The mortality associated with SARS-CoV is substantially higher than that associated with SARS-CoV-2 (www.who.int) [Dong et al., 2020; Nishiura et al., 2020a; Pan et al., 2020; Rothe et al., 2020]. While SARS-CoV causes a disease of the lower respiratory tract, and infected individuals are only contagious when they experience severe symptoms [de Wit et al., 2016], SARS-CoV-2 is present in the upper respiratory tract and seems to be readily transmitted via droplets and direct contact prior to the onset of symptoms. Moreover, mild but infectious cases may substantially contribute to its spread [Li et al., 2020; Rothe et al., 2020; Yang et al., 2020; Yu et al., 2020]. Further research will be required to elucidate in detail, which DCPs are responsible for which differences in virus behaviour.

However, we have already identified a number of differences between SARS-CoV-2 and SARS-CoV, with regard to their cell tropism and drug sensitivity profiles. Both viruses use ACE2 as a receptor and are activated by the transmembrane serine protease TMPRSS2 [Li et al., 2003; Matsuyama et al., 2010; Cui et al., 2019; Hoffmann et al., 2020; Letko et al., 2020; Lu et al., 2020; Matsuyama et al., 2020; Wals et al., 2020; Wan et al., 2020; Wetko et al., 2020; Wrapp et al., 2020; Yan et al., 2020; Zhou et al., 2020]. Our results show, however, that the ACE2 and the TMPRSS2 status are not sufficient to predict cells susceptibility to SARS-CoV-2 or SARS-CoV. We found that the colorectal cancer cell line CL14 supported SARS-CoV-2 replication, although it displayed lower ACE2 levels and similar TMPRSS2 levels to the non-susceptible cell lines DLD-1 and HT29. Hence CL14 represents a novel additional model for the studying of SARS-CoV-2 replication. Notably, attempts to identify SARS-CoV-2 target cells based on the ACE2 status [Luan et al., 2020; Qiu et al., 2020; Xu et al., 2020] need to be considered with caution in the light of our current findings.

As previously described [Kamitani et al., 2006], ACE2 expression rendered SARS-CoV non-permissive 293 cells susceptible to SARS-CoV. However, the effects of ACE2 expression had a substantially lower impact on SARS-CoV-2 infection, indicating differences in other host cell determinants of SARS-CoV and SARS-CoV-2 susceptibility. Moreover, an anti-ACE2 antibody displayed higher efficacy against SARS-CoV than against SARS-CoV-2. This may be explained by an increased SARS-CoV-2 S affinity to ACE2 compared to SARS-CoV S [Wrapp et al., 2020], which may be more difficult to antagonise.

The serine protease inhibitors camostat, nafamostat, and aprotinin inhibited both SARS-CoV-2 and SARS-CoV CPE formation. In contrast to aprotinin, camostat, and nafamostat exerted limited activity against double-stranded RNA formation in SARS-CoV-2-infected cells. This may indicate that camostat and nafamostat rather exert cytoprotective effects that prevent cells from virus-induced lysis but less pronounced antiviral activity. The mechanisms underlying the enhanced anti-SARS-CoV-2 activity of aprotinin remain unclear. Differences in the interference with additional proteases involved in SARS-CoV-2 replication may be responsible. Notably, aprotinin had been identified in the past as a protease inhibitor with pronounced antiviral activity, which may interfere with viral proteases in addition to cellular ones [Hayashi et al., 1991; Aleshin et al., 2007; Zhirnov et al., 2011; Lin et al., 2017].

Notably, SARS-CoV-2 was more sensitive to aprotinin than SARS-CoV, which may be at least in part explained by the DCPs observed in the vicinity of the cleavage sites in S. Effective aprotinin concentrations were in the range of clinically achievable concentrations. Moreover, aprotinin aerosols, which may result in increased local drug concentrations in the lungs are approved for the treatment of influenza viruses in Russia [Zhirnov et al., 2011]. Remdesivir, a broad spectrum antiviral agent under investigation in clinical trials for the treatment of SARS-CoV-2 patients (www.clinicaltrials.gov), exerted stronger effects against SARS-CoV than against SARS-CoV-2.

Therapeutic concentrations of the proton pump inhibitor omeprazole further increased the activity of aprotinin and remdesivir. Omeprazole may interfere with the acidification of the lysosomes, which is required to support coronavirus replication [Shen et al., 2017]. However, other, so far unknown, mechanisms may also contribute to this. Notably, omeprazole and other proton pump inhibitors have recently been shown to increase the anti-herpes simplex virus activity of acyclovir [Michaelis et al., 2019].

In conclusion, our in-silico study revealed a substantial number of differentially conserved amino acid positions in the SARS-CoV-2 and SARS-CoV proteins. In agreement, cell culture experiments identified differences in the cell tropism and drug sensitivity profiles of these two viruses. Our data also show that cellular ACE2 levels do not reliably indicate cell susceptibility to SARS-CoV-2. Hence, ACE2 expression studies are not sufficient to predict the SARS-CoV-2 cell tropism. Differences in the drug sensitivity profiles between SARS-CoV-2 and SARS-CoV, the most closely related coronavirus known to have caused disease in humans, indicate that approaches to identify anti-SARS-CoV-2 drugs will require testing against this virus. Finally, and probably most importantly during an ongoing pandemic, we have shown that the approved drug aprotinin inhibits SARS-CoV-2 infection in clinically achievable concentrations. The efficacy of aprotinin (and of remdesivir, which is investigated against SARS-CoV-2 in clinical trials) can be further enhanced by therapeutic concentrations of the proton pump inhibitor omeprazole. Hence, our study has identified novel candidate therapies based on approved drugs that can be readily tested in a clinical setting.

## Methods

### Structural analysis

Full genome sequences for SARS-CoV-2 were obtained from the National Center for Biotechnology Information (NCBI) 4 and the GISAID resource. A total of 1266 full length genome sequences were available as of 27/03/2020. Fifty-three SARS-CoV genome sequences were downloaded from VIPR [Pickett et al., 2012; Pickett et al., 2012A] restricted to sequences with a collection year between 2003-2004 and a human or unknown host. Where the host was unknown the genome information was further checked to see if it was appropriate. Open Reading Frames (ORFs) were extracted using EMBOSS getorf [Rice et al., 2000]. These ORFs were matched to known proteins using BLAST, and fragments and mismatches were discarded. To match the ORF1ab non-structural proteins, a BLAST database of the sequences from the SARS non-structural proteins was generated and the SARS-CoV-2 ORF1ab searched against it. After each ORF was assigned to a known protein they were aligned using ClustalO [Sievers et al., 2011] with default settings. Sequences that fell below 50% coverage were removed from analysis.

SDPs were identified by calculating the Jensen-Shannon divergence [Capra & Singh, 2007] score for each position in the multiple sequence alignment in each species. Highly conserved alignment positions where the conservation score was >0.8 for both species were retained. Any of alignment positions where the same amino acid occurred in both species were then removed. The remaining residues, were considered SDPs.

All available SARS-CoV-2 and SARS-CoV protein structures were downloaded from the protein Databank (PDB) [Armstrong et al., 2020]. Where structures were not available they were modelled using Phyre2 [Kelley et al., 2015] (Supplementary Table 4). SDPs were mapped onto protein structures using PyMOL from structures obtained from the Protein Databank (PDB). To model the complex between the SARS-CoV-2 spike protein and ACE2, a model of the SARS-CoV-2 spike protein was built using Phyre2 based on the SARS-CoV structure (PDB:6acg) as some of the residues involved in binding by the SARS-CoV spike protein were not resolved in the SARS-CoV-2 spike protein structure. This homology model was docked to ACE2 using HADDOCK with constraints based on the likely interface residues equivalent to the SARS-CoV complex.

### Cell culture

The Caco2 cell line was obtained from DSMZ (Braunschweig, Germany). The cells were grown at 37°C in minimal essential medium (MEM) supplemented with 10% foetal bovine serum (FBS), 100 IU/ml penicillin, and 100 μg/ml of streptomycin. 293 cells (PD-02-01; Microbix Bisosystems Inc.) and 293/ACE2 cells [Kamitani et al., 2006] (kindly provided by Shinji Makino, UTMB, Galveston, Texas) were cultured in Dulbecco’s modified Eagle medium (DMEM) supplemented with 10% FBS), 50 IU/ mL penicillin, and 50µg/ mL streptomycin. Selection of 293/ACE2 cells constitutively expressing human angiotensin-converting enzyme 2 (ACE2) was performed by addition of 12 µg/ mL blasticidin. All culture reagents were purchased from Sigma (Munich, Germany). Cells were regularly authenticated by short tandem repeat (STR) analysis and tested for mycoplasma contamination.

### Virus infection

The isolate SARS-CoV-2/1/Human/2020/Frankfurt was derived from an individual, who had been evacuated from the Hubei province in China, transferred to University Hospital Frankfurt, and tested positive for SARS-CoV [Hoehl et al., 2020] and cultivated in Caco2 cells as previously described for SARS-CoV strain FFM-1 [Cinatl et al., 2004]. Virus titres were determined as TCID50/ml in confluent cells in 96-well microtitre plates [Cinatl et al., 2003; Cinatl et al., 2005].

### Western blot

Cells were lysed using Triton-X-100 sample buffer, and proteins were separated by SDS-PAGE. Detection occurred by using specific antibodies against β-actin (1:2500 dilution, Sigma-Aldrich, Munich, Germany), ACE2, and TMPRSS2 (both 1:1000 dilution, abcam, Cambridge, UK). Protein bands were visualised by laser-induced fluorescence using infrared scanner for protein quantification (Odyssey, Li-Cor Biosciences, Lincoln, NE, USA).

### Receptor blocking experiments

To investigate whether ACE2 or DPP4 receptors are involved in SARS-CoV-2 internalisation and replication, Caco2 cells were pre-treated for 30 min at 37°C with goat antibody directed against the human ACE2 or DDP4 ectodomain (R&D Systems, Wiesbaden-Nordenstadt, Germany). Then, cells were washed three times with PBS and infected with SARS-CoV-2 at MOI 0.01. Cytopathogenic effects were monitored 48h post infection.

### Antiviral assay

Confluent cell cultures were infected with SARS-CoV-2 in 96-well plates at MOI 0.01 in the absence or presence of drug. Cytopathogenic effect (CPE) was assessed visually 48h post infection [Cinatl et al., 2003].

### Viability assay

Cell viability was determined by 3-(4,5-dimethylthiazol-2-yl)-2,5-diphenyltetrazolium bromide (MTT) assay modified after Mosman [Mosmann, 1983], as previously described [Onafuye et al., 2019]. Confluent cell cultures in 96-well plates were incubated with drug for 48h. Then, 25 µL of MTT solution (2 mg/mL (w/v) in PBS) were added per well, and the plates were incubated at 37 °C for an additional 4 h. After this, the cells were lysed using 200 µL of a buffer containing 20% (w/v) sodium dodecylsulfate and 50% (v/v) N,N-dimethylformamide with the pH adjusted to 4.7 at 37 °C for 4 h. Absorbance was determined at 570 nm for each well using a 96-well multiscanner. After subtracting of the background absorption, the results are expressed as percentage viability relative to control cultures which received no drug. Drug concentrations that inhibited cell viability by 50% (IC50) were determined using CalcuSyn (Biosoft, Cambridge, UK).

### qPCR

SARS-CoV-2 and SARS-CoV RNA from cell culture supernatant samples was isolated using AVL buffer and the QIAamp Viral RNA Kit (Qiagen) according to the manufacturer’s instructions.

Absorbance-based quantification of the RNA yield was performed using the Genesys 10S UV-Vis Spectrophotometer (Thermo Scientific). RNA was subjected to OneStep qRT-PCR analysis using the Luna Universal One-Step RT-qPCR Kit (New England Biolabs) and a CFX96 Real-Time System, C1000 Touch Thermal Cycler. Primers were adapted from the WHO protocol29 targeting the open reading frame for RNA-dependent RNA polymerase (RdRp): RdRP_SARSr-F2 (GTG ARA TGG TCA TGT GTG GCG G) and RdRP_SARSr-R1 (CAR ATG TTA AAS ACA CTA TTA GCA TA) using 0.4 µM per reaction. Standard curves were created using plasmid DNA (pEX-A128-RdRP) harbouring the corresponding amplicon regions for RdRP target sequence according to GenBank Accession number NC_045512. For each condition three biological replicates were used. Mean and standard deviation were calculated for each group.

### Immunostaining for double-stranded RNA

Immunostaining was performed as previously described [Cinatl et al., 1995], using a monoclonal antibody directed against double-stranded RNA (1:150 dilution, SCICONS J2, mouse, IgG2a, kappa chain, English & Scientific Consulting Kft., Szirák, Hungary) 48h post infection.

## Acknowledgements

The authors thank Shinji Makino, UTMB, Galveston, TX, for the provision of 293/ACE2 cells.

## Data availability

All data are provided in the manuscript and its supplements.

## Author contributions

D.B., J.E.M, K.M., S.G.M., M.N.W., and J.C. performed experiments. V.K. provided essential materials. All authors analysed data. M.N.W., M.M., and J.C. planned, conducted, and supervised the study. M.M. wrote the first manuscript draft. All authors were involved in the drafting of and approved the final manuscript version.

