## Supplementary material for "SARS-CoV-2 and SARS-CoV differ in their cell tropism and drug sensitivity profiles": Suppl Material

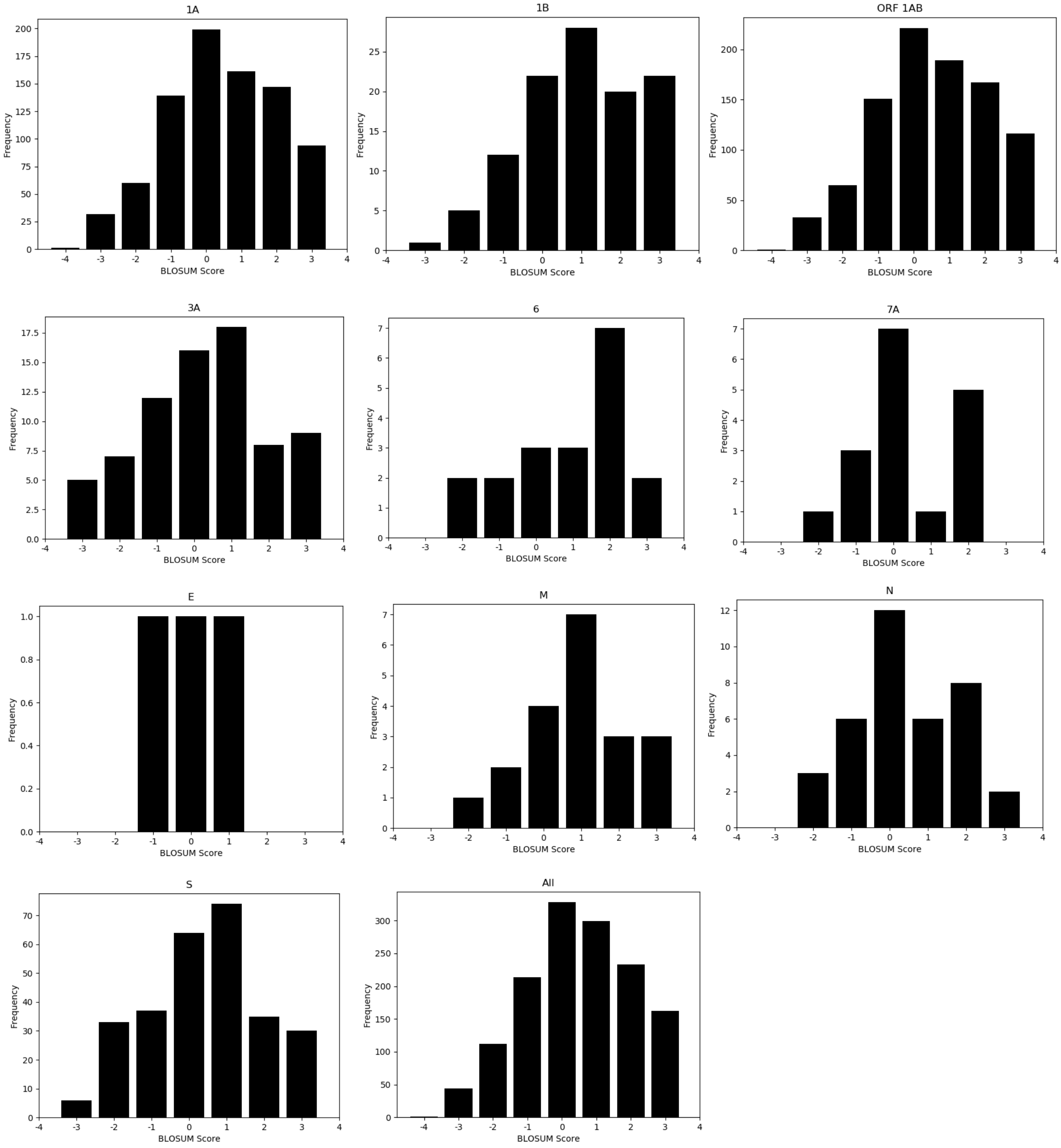


**Supplementary Figure 1.** The BLOSUM scores for the amino acid substitutions present in the SDPs. A graph is plotted that combines all of the proteins and one for each of the individual proteins that were analysed.

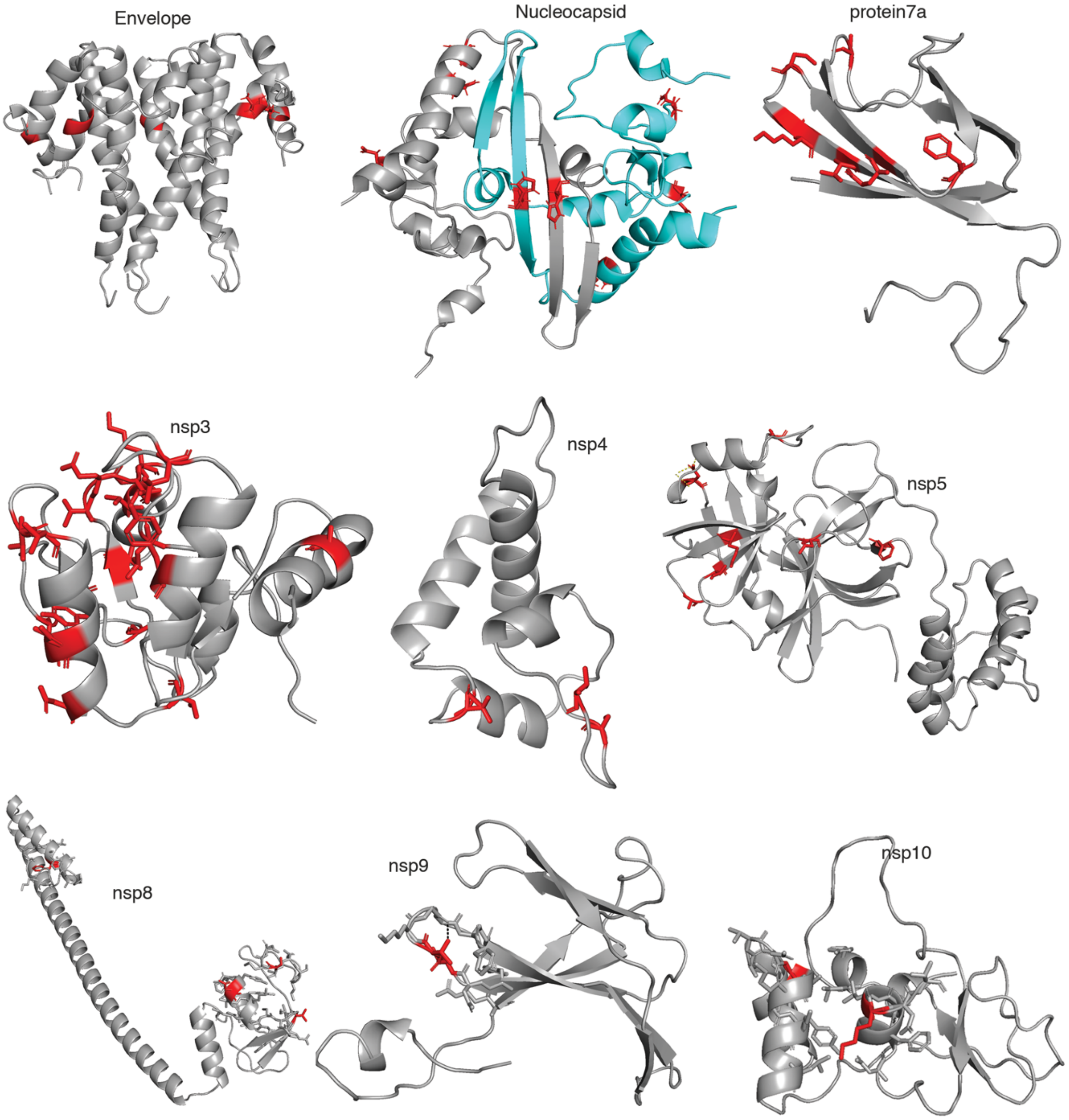


**Supplementary Figure 2.** Overview of modelled DCPs.

**Supplementary Table 2 - Analysis of DCPs present in the SARS-CoV and SARS-CoV-2 Spike protein interface with human ACE2.**

| SDP | SARS-CoV structural analysis | SARS-CoV-2 structural analysis | Effect? |
|---|---|---|---|
| V404=K417 | V404 is not in the interface | K417 is in the interface and could form a salt bridge with ACE-2 D30 | Likely – new polar interaction within interface |
| R426=N439 | Loss of hydrogen bond to ACE2 Gln325 due to shorter sidechain. N would still be able to form hydrogen bonds | N439 is located away from the interface site and so does not form a hydrogen bond with ACE2. Instead forms a hydrogen bond with S443 (also a DCP – A430=S443) which is likely to stabilise the loop they are both part of. | Likely – Loss of interface hydrogen bond. |
| Y442=L455 | Y422 forms hydrogen bond to backbone of W476 – loss could result in conformational change. The sidechain also contacts the backbone of ACE2 D30 and K31 | L455 remains in interface and contacts ACE2 D30 and H34. | Likely – loss of intramolecular hydrogen bond |
| F460=Y473 | Conservative change. | Introduction of OH group that can form hydrogen bonds. Y473 forms hydrogen bond with backbone of R457 and is closer to ACE2 T27 so potential to form hydrogen bond in interface. | Possible – introduction of hydrogen bond (could be with ACE2) |
| P462=A475 | Located in a loop, could affect this conformation – many DCPs in this loop | Loop has different conformation. | Possible – Conformational change of loop |
| N479=Q493 | Interface hydrogen bond formed with ACE2 H34 backbone. With a shorter sidechain this this may be lost in SARS-CoV-2. | Q493 forms a hydrogen bond with ACE2 E35 in this complex. So hydrogen bond is maintained but also different. | Possible – hydrogen bond with ACE2 retained but to different residue. |
| Y484=Q498 | Y484 can form hydrogen bonds with ACE2 Gln42 (sidechain) and intramolecular H bonds with T433 (backbone), Y436 (sidechain). | Q498 maintains hydrogen bonds with ACE2 Gln42 | Possible – change in residue forming hydrogen bonds with ACE2. |
| T485=P499 | Sidechain points away from interface, loss of hydrogen bond with R426 (also a | Loop conformation similar as for SARS-CoV structure but not coordinated with other loop | Likely - loss of intramolecular hydrogen bond |

| | | | |
|---|---|---|---|
| | DCP) backbone in adjacent loop. This hydrogen bond is likely to coordinate the structure between these two loops. There are multiple DCPs present in both loops | | |
| I489=V503 | Conservative change I489 in direct contact with ACE2 Q325 | slightly smaller sidechain is further away from ACE2 Q325. | Unlikely. |
